# Stress Recovery Through Mindful Breathing: Convergent Neural Signatures in Around-Ear and Scalp EEG

**DOI:** 10.64898/2026.04.21.719740

**Authors:** Stephanie Nelli, Pachaya Sailamul, Waraset Wongsawat, Suparach Intarasopa, Tanagrit Phangwiwat, Achirawich Sombatsompop, Matsui Hiroshi, Sarawin Khemmachotikun, Chitpol Mungprom, Barry Giesbrecht, Sirawaj Itthipuripat

## Abstract

Mindfulness-based interventions effectively reduce stress and anxiety, yet the neural mechanisms underlying contemplative practices and their optimal implementation parameters remain poorly understood. A critical barrier to real-world application is the absence of validated minimally invasive neural recording technologies. Here, we simultaneously recorded full-coverage scalp and around-ear EEG during a 2×2×2 factorial design manipulating (1) cognitive state (mental arithmetic stress vs. passive viewing), (2) recovery strategy (mindful breathing meditation vs. mind-wandering), and (3) sensory context (eyes open vs. eyes closed). Mental arithmetic robustly elevated subjective stress and modulated canonical oscillatory patterns: increased midline frontal theta power (3-7 Hz), suppressed posterior alpha power (10-12 Hz), and enhanced posterior beta and gamma power (25-48 Hz). All rest conditions reduced subjective stress following stress induction, with eyes-closed mindful breathing producing maximal reduction. Critically, mindful breathing differentially modulated temporal beta and gamma power in a context-dependent manner, with effects determined by prior cognitive state and eye position. Eyes-closed meditation maximally suppressed gamma power within 20 seconds following arithmetic stress, whereas eyes-open meditation alone was sufficient for gamma suppression following passive viewing. Around-ear electrodes detected these stress and meditation signatures with comparable fidelity to scalp recordings. These findings reveal that mindful breathing engages rapid, context-dependent neural regulation mechanisms and establish that wearable EEG can reliably capture these dynamics, enabling real-world stress monitoring and mindfulness guidance.

**Impact Statement:** Mindfulness-based interventions reduce stress, yet their neural mechanisms and optimal implementation remain unclear, partly due to limited real-world neural measurement tools. Using simultaneous scalp and around-ear EEG, we show that mindful breathing rapidly and context-dependently regulates stress-related brain activity via changes in high-frequency EEG oscillations (i.e., gamma band activity), with effects shaped by prior cognitive state and eye condition. Importantly, around-ear EEG captured these neural signatures with fidelity comparable to scalp recordings, enabling wearable neurotechnology for real-world stress monitoring and personalized mindfulness guidance.

## Introduction

Mindfulness meditation, particularly focused attention on breathing, constitutes a core component of evidence-based interventions for stress reduction, anxiety management, and mood regulation (Goyal et al., 2014; Grossman et al., 2004; Kabat-Zinn, 2009; Kabat-Zinn et al., 1985; Segal et al., 2002; Tang et al., 2015). Despite robust clinical efficacy and extensive neuroscientific investigation, critical gaps remain in understanding how specific practice parameters modulate neural mechanisms and how laboratory findings translate to real-world applications.

EEG studies have identified distinct neural correlates of mindfulness meditation across multiple frequency bands, though mechanistic interpretations remain debated. Frontal midline theta oscillations (∼3-7 Hz) increase during focused attention meditation, likely reflecting sustained cognitive control and performance monitoring (Kubota et al., 2001; Lagopoulos et al., 2009; Lee et al., 2018; Tang et al., 2019; cf. Cavanagh & Frank 2014; Itthipuripat et al., 2013, 2019; Lomas et al., 2015; Rungratsameetaweemana et al., 2018). Alpha oscillations (∼10-12 Hz) exhibit complex, state-dependent modulation. During meditation with eyes closed, posterior alpha power typically increases, consistent with enhanced cortical inhibition or reduced sensory processing (Barry et al., 2007; Berger, 1929; Cahn & Polich, 2006; Jensen & Mazaheri, 2010; Toscani et al., 2010). Conversely, alpha suppression accompanies heightened attentional engagement during demanding cognitive tasks (Banerjee et al., 2011; Bonnefond & Jensen, 2025; Foxe & Snyder, 2011; Itthipuripat et al., 2019, 2023; Klimesch et al., 1998, 2007, 2012; Nelli et al., 2017; Sookprao, 2024; Strauß et al., 2014; Woodman et al., 2022). High-beta (∼20-32 Hz) and gamma (∼35-100 Hz) oscillations in frontal, parietal, and temporal cortex have been associated with cognitive integration, sustained attention, and expert meditation practice (Berkovich-Ohana, et al., 2012; Cahn, et al., 2010; Huang and Lo, 2009; Kaur and Singh, 2015; Lehmann et al., 2001; Lutz et al., 2004; Vialatte et al., 2019), potentially reflecting excitatory-inhibitory balance in cortical circuits (Buzsáki & Wang, 2012; Cardin, 2016; Sohal & Rubenstein, 2019; Wang & Buzsaki, 1996). However, considerable heterogeneity exists across studies due to varied meditation traditions, task instructions, and recording methodologies (Davidson & Kaszniak 2015).

A fundamental unresolved question concerns how specific practice parameters modulate neural stress responses. Eye state represents a particularly important yet understudied parameter. Eye closure substantially reduces visual input and increases posterior alpha power (Barry et al., 2007; Berger, 1929; Cahn & Polich, 2006; Jensen & Mazaheri, 2010; Toscani et al., 2010), yet whether this sensory manipulation interacts with meditation to enhance neural stress regulation remains unknown. Critically, if eye closure enhances meditation efficacy, interventions could be optimized by matching practice instructions to environmental contexts. The temporal dynamics of acute meditation effects remain poorly characterized. Longitudinal studies document gradual neural plasticity over weeks to months of training (Cahn et al., 2006; Hölzel et al., 2011; Itthipuripat et al., 2017; Saggar et al., 2012; Skwara et al., 2012; Yoshida et al., 2020), but the timescale at which acute meditation practice modulates neural activity within a single session remains largely unexplored. Understanding whether effects emerge rapidly at practice onset versus accumulating gradually would inform the design of brief, accessible interventions suitable for real-world implementation.

A second critical gap concerns validating wearable neurotechnology for capturing meditation-relevant neural dynamics in naturalistic settings. Conventional full-coverage scalp EEG is impractical for ambulatory monitoring due to extensive preparation requirements, physical discomfort, and poor aesthetic acceptability. Around-ear EEG systems offer a minimally invasive alternative that enables continuous neural monitoring during daily activities (Athavipach et al., 2019; Bleichner & Debener, 2017; Bleichner et al., 2015; Debener et al, 2015; Hemakom et al., 2024; Israsena & Pan-ngum, 2022; Kaongoen et al., 2021; Sanguantrakul et al., 2024;, see reviews Choi et al., 2023; Correia et al., 2024; da Silva Souto et al., 2022; Geirnaert et al., 2025; Hans et al., 2022; Kaongoen et al., 2023; Knierim et al., 2023; Mai et al., 2025; Petrossian et al., 2023). While around-ear systems reliably capture evoked potentials and tonic oscillatory rhythms, their sensitivity to dynamic neural state changes relevant for mental health applications remains unvalidated. Critically, around-ear placements provide limited spatial coverage relative to full-scalp montages, raising questions about which stress and meditation-related neural signatures can be reliably detected.

The present study addressed these gaps through a within-subjects design combining mental arithmetic stress induction with subsequent rest periods involving either mindful breathing meditation or mind-wandering, performed with eyes open or closed. We simultaneously recorded full-coverage scalp EEG and around-ear EEG to directly compare signal quality and spatial sensitivity across recording methods. We hypothesized that mental arithmetic would elicit canonical stress-related neural responses including increased frontal theta, suppressed posterior alpha, and elevated posterior and temporal beta/gamma activity, consistent with prior literature (Cavanagh & Frank, 2014; Fernández et al., 1995; Foxe & Snyder, 2011; Gashaj et al, 2024; Gruber et al., 1999; Ishii et al., 2014; Itthipuripat et al., 2013, 2019, 2023; Katahira, et al., 2018; Katsumi, et al., 2023; Klimesch et al., 1998, 2007, 2012; Müller, et al., 2000; Nelli et al., 2017; Rungratsameetaweemana et al., 2018; Sakkalis et al, 2006; Simos et al, 2002; Sookprao, 2024; Williamson et al., 1997; Woodman et al., 2022). We predicted that mindful breathing would reduce subjective stress and modulate stress-related oscillatory activity, particularly high-beta and gamma oscillations reflecting cortical excitatory-inhibitory balance (c.f., Attar et al, 2022; Buzsáki & Wang, 2012; Choi, et al., 2015; Ehrhardt et al., 2021; Hayashi, et al., 2009; Jena et al., 2015; Palacios-Gracia, et al., 2021; Rusinova et al., 2024; Seo, et al., 2010; Snijders et al., 2013; Sohal & Rubenstein, 2019; Travis & Shear, 2010; Yokoyama et al., 2025). We hypothesized that meditation effects would vary systematically with eye state, revealing context-dependent neural mechanisms of contemplative stress regulation.

Finally, we assessed whether around-ear EEG could capture stress and meditation signatures with fidelity comparable to scalp recordings. We predicted that around-ear electrodes would reliably detect temporal cortical activity due to favorable anatomical proximity, but would show reduced sensitivity to posterior and frontal sources given limited spatial coverage. Establishing the capabilities and limitations of around-ear EEG for stress and meditation monitoring would provide essential validation for translating laboratory findings into practical, ambulatory neurofeedback applications that could enhance mindfulness intervention accessibility and personalization.

## Methods and Materials

### Subjects

We recruited 35 neurologically and psychologically healthy adults (age range = 18–36 years, M = 22.08, SD = 3.73; 13 female; 33 right-handed), with normal or corrected-to-normal vision and hearing. Written informed consent was obtained from all participants prior to their participation in the study, and all procedures were conducted in accordance with the Declaration of Helsinki and approved by the Institutional Review Board of King Mongkut’s University of Technology Thonburi. Five participants were excluded due to technical recording errors resulting in missing event triggers, yielding a final sample of 30 participants (age range = 18–36 years, M = 22.17, SD = 3.94; 9 females; 29 right-handed).

### Stimuli and Task

Stimulus presentation was programmed in MATLAB version 2020b (MathWorks 2020) using Psychtoolbox (Brainard 1997; Pelli 1997). Visual stimuli were displayed on a gray background (luminance = 300 cd/m²) on an LED monitor (refresh rate = 165 Hz) controlled by a personal computer with the Windows 10 operating system. Participants were instructed to maintain central fixation throughout all blocks, minimize eye and head movements, and remain as still as possible to reduce EEG artifacts. The display monitor was positioned on an adjustable standing desk, allowing the fixation cross to be aligned vertically with each participant’s eye level in both sitting and standing postures.

The experiment comprised eight block types in a 2×2×2 factorial design: task context (arithmetic vs. passive viewing) × rest condition (mindful breathing vs. mind-wandering) × eye state (open vs. closed). Block order was counterbalanced across participants to minimize order effects. Half of the participants performed eight blocks while seated followed by eight blocks while standing; the remaining half completed the posture order in reverse. Each block began with an instruction screen indicating whether the upcoming task would involve arithmetic calculation or passive viewing. Participants then provided baseline ratings of subjective stress, arousal, and concentration using a 0–9 Likert scale (0 = extremely low; 5 = moderate; 9 = extremely high), responding as accurately as possible.

Participants then completed six difficult arithmetic problems (multiplication or division). To ensure adequate and comparable cognitive load, multiplication problems consisted of a three-digit number (randomly drawn from 121–149) multiplied by a one-digit number (randomly drawn from 3–4 and 6–9), whereas division problems used the same numerical structure in inverse form, with the resulting three– to four-digit products divided by the corresponding one-digit number (randomly drawn from 3–4 and 6–9). Each problem was displayed for 15 seconds with an on-screen countdown (from 15 to 0 s). Participants stated their answers aloud while a research assistant present in the room recorded responses, introducing an element of social-evaluative stress. After each 15-second viewing interval, a 5-second response window appeared, during which participants entered their answers using a keyboard (digits 0–9) with both hands. In passive viewing blocks, participants viewed visually identical math problems but were instructed not to compute answers. Instead, they verbally indicated whether each problem involved multiplication or division during the 15-second viewing window, and then pressed the corresponding keyboard button during the 5-second response prompt.

Following the arithmetic calculation or passive viewing phase, participants completed a second set of subjective ratings (stress, arousal, concentration; 5 seconds per scale). Participants then entered a 120-second rest period in which they were instructed either to allow their thoughts to unfold freely (i.e., mind-wandering) or to focus on their breathing and bodily sensations (i.e., meditation), with their eyes open or closed. A bell tone signaled the end of the rest period, cueing participants to stop meditating (and open their eyes if closed) and complete a final rating sequence. Each block lasted approximately 5 minutes, followed by a 2–3-minute break. Including EEG setup, task instructions, and all 16 blocks, total participation time was approximately 3-3.5 hours.

### Behavioral Analysis

We first averaged participants’ subjective ratings of arousal, stress, and concentration across subjects at each time point (baseline, post-task, and post-rest). Within-subject standard errors of the mean (SEM) were computed for all measures.

To examine the effects of task context and time period, we conducted separate 2×3 repeated-measures ANOVAs with task context (arithmetic vs. passive viewing) and period (baseline, post-task, post-rest) as within-subject factors for arousal, stress, and concentration scores. We then conducted planned paired t-tests to clarify effects in three steps. First, we compared baseline ratings between the arithmetic and passive viewing blocks at the baseline. Similar analyses were conducted on the post-task and post-rest ratings to assess whether elevated levels of stress, arousal, and concentration occurred and persisted throughout the block. Second, we compared post-task ratings against baseline ratings for each task context separately to assess the effectiveness of the arithmetic task in elevating arousal, stress, and concentration relative to passive viewing. Third, we compared post-rest ratings with both baseline and post-task ratings, again separately for the arithmetic and passive viewing conditions, to evaluate recovery dynamics and determine whether reductions in arousal, stress, and concentration returned to or remained elevated above baseline levels. Together, these analyses tested whether stress, arousal, and concentration increased after the arithmetic task, whether passive viewing produced minimal change, and whether the 2-minute resting period, with meditation or mind-wandering, reduced subjective stress, arousal, and concentration.

Next, we evaluated how meditation (mindful breathing vs. mind-wandering), eye state (eyes closed vs. open), and task context (post-arithmetic calculation vs. post-passive viewing), as well as their interactions, influenced subjective ratings by running repeated-measures ANOVAs on scores collected during the post-rest period. Significant interaction effects were followed up with post hoc paired t-tests across combinations of meditation and eye-state conditions.

### EEG recording, preprocessing, and analysis

Continuous EEG was recorded using a 64-channel ActiveTwo system (BioSemi, Amsterdam, Netherlands) with eight additional external electrodes, sampled at 512 Hz. The external electrodes consisted of bilateral paired placements on the mastoids (used as references), 2.5 cm inferior to each mastoid behind the ears, at the left and right temples near the upper front of the ears, and at the anterior opening of the left and right ear canals. EEG data were recorded online referenced to the common mode sense–driven right leg (CMS–DRL) electrode. All channel offsets were kept below 20 mV, consistent with standard operation guidelines for the BioSemi ActiveTwo system.

Data preprocessing and analysis were performed using EEGLAB v2020.0 (Delorme & Makeig, 2004) and custom MATLAB scripts. Continuous EEG signals were rereferenced offline to the average of the left and right mastoid electrodes, bandpass filtered between 0.25 and 100 Hz using a third-order Butterworth filter, and notch filtered from 49–51 Hz to remove line noise. Independent component analysis (ICA) was then performed to identify and remove components reflecting eye blinks, ocular and head movements, muscle activity, slow drifts, and other non-neural artifacts (Makeig et al., 1996).

To examine oscillatory power modulations in both scalp and around-ear EEG during stress induction across the arithmetic and passive viewing conditions, we segmented the artifact-corrected continuous EEG into six consecutive 20-s bins spanning 0-120 s relative to task onset (0-20, 20-40, 40-60, 60-80, 80-100, and 100-120 s).For each bin, block, electrode, and participant, we computed power spectral density (PSD) estimates (1-48 Hz) using Welch’s method (i.e., pwelch MATLAB built-in; Welch, 1967). Power values were log-transformed to decibels and averaged across the six temporal bins. This temporal binning and subsequent averaging were implemented to minimize the influence of residual artifacts that may remain in the continuous data. For the scalp PSD data, we averaged electrodes to form eight spatial regions using arithmetic means, including the midline frontal (F1, Fz, F2), left and right central (C1, C3 and C2, C4), left and right temporal (C5, T7 and C6, T8), left and right posterior occipital (P9, PO7 and PO8, P10), and the mid-occipital region (O1, Oz, O2). For the around-ear EEG data, we analyzed all six electrode locations independently, including the left and right positions in front of the ear, at the temple, and behind the ear. To visualize the group-level results, we applied an additional moving average (using MATLAB’s movmean function) to smooth the power spectral density (PSD) estimates. We then computed grand-average PSDs across all participants along with within-subject standard errors of the mean (SEM; Cousineau, 2005; Morey, 2008) Importantly, this smoothing procedure was used only for plotting purposes and all statistical analyses were performed on the non-smoothed data.

For scalp EEG, we conducted repeated-measures ANOVAs to test main effects of task context and posture, as well as their interaction, on four canonical oscillatory bands modulated by cognitive stress. We examined: (i) midline frontal theta power (3–8 Hz), associated with cognitive control, performance monitoring, and working memory maintenance (Cavanagh & Frank, 2014; Gärtner et al., 2015; Itthipuripat et al., 2013); (ii) posterior alpha power (8–12 Hz), reflecting attentional disengagement and cortical inhibition, and thus predicted to decrease during arithmetic (Banerjee et al., 2011; Bonnefond & Jensen, 2025; Foxe & Snyder, 2011; Itthipuripat et al., 2019, 2023; Jensen & Mazaheri, 2010; Klimesch et al., 1998, 2007, 2012; Nelli et al., 2017; Sookprao, 2024; Strauß et al., 2014; Woodman et al., 2022); (iii) central beta power (13–25 Hz), which typically decreases during motor preparation and execution (Kilavik et al., 2013; Pfurtscheller & Lopes da Silva, 1999); and (iv) temporal and posterior high-beta (26–33 Hz) and gamma (35–48 Hz) power, reflecting enhanced excitatory drive during visual processing, stress and arousa l(Attar et al, 2022; Buzsáki & Wang, 2012; Choi, et al., 2015; Ehrhardt et al., 2021; Hayashi, et al., 2009; Jena et al., 2015; Palacios-Gracia, et al., 2021; Ray & Maunsell, 2011; Seo, et al., 2010; Snijders et al., 2013; Sohal & Rubenstein, 2019; Travis & Shear, 2010; Yokoyama et al., 2025).

For around-ear EEG, we applied the same statistical framework to theta, alpha, low-beta, high-beta, and gamma band activity at each of the six electrode contacts. Multiple comparisons across these recording sites were controlled using Holm–Bonferroni correction (Holm, 1979).

For the rest-period data, we applied the same time binning and spectral decomposition procedures used during the stress-induction period. Specifically, artifact-corrected continuous EEG was segmented into time bins and transformed using Welch’s method, and power values were converted to decibels prior to averaging. First, we examined whether task context and posture modulated key oscillatory components in a manner similar to those observed during the stress-induction period. To this end, we performed repeated-measures ANOVAs with task context and posture as within-subject factors on both scalp EEG and around-ear EEG data. Oscillatory powers were collapsed across all time bins in the same manner as in the analyses conducted for the stress-induction period.

To characterize the temporal dynamics of meditation– and eye-state–related effects, we divided the rest period into two windows: (i) the initial 0–20 s following rest onset and (ii) the sustained 20–120 s segment. Note that the latter window collapsed across 20–40, 40–60, …, 100–120 s bins because oscillatory patterns were highly consistent across these intervals.

We then performed repeated-measures ANOVAs to test the effects of meditation style (mindful breathing vs. mind-wandering), eye state (eyes closed vs. open), task context (post-arithmetic vs. post-passive viewing), and time window (0-20 s vs. 20-120 s), as well as their interactions, on midline frontal theta activity, posterior occipital alpha activity, motor-related low-beta activity over central scalp sites, and high-beta and gamma activity at temporal and posterior occipital regions. For each temporal window, we ran separate repeated-measures ANOVAs to examine the main effects of meditation, eye state, and task context and their interactions on posterior occipital alpha power, where robust modulation by eye state was observed. Because the main effects of eye state on alpha-band activity were observed across the entire scalp as well as in the around-ear electrodes, we additionally performed statistical analyses at all scalp and around-ear electrode positions. Multiple comparisons were corrected using the Holm–Bonferroni method. Parallel analyses were conducted on gamma-band activity recorded over bilateral temporal scalp regions, where modulation was driven by task context, meditation style, and eye state. As we observed a significant interaction among meditation style, eye state, and task context on gamma-band activity in the bilateral temporal regions, we conducted post hoc paired t-tests within each task context to compare eye-open meditation versus eye-open mind-wandering and eye-closed meditation versus eye-closed mind-wandering, separately for the 0–20 s and 20–120 s windows.

For the around-ear EEG, we focused statistical analyses on alpha– and gamma-band activity, corresponding to the effects observed in scalp EEG (eye-state modulation for alpha and task-context modulation for gamma). As with scalp data, we used repeated-measures ANOVAs to examine the effects of meditation style, eye state, task context, and time window, as well as their interactions, on alpha and gamma power across around-ear contacts. When significant main effects or interactions were identified, we followed the same procedure used for scalp temporal gamma activity, conducting separate ANOVAs and post hoc comparisons to characterize condition-wise differences using paired t tests.

## Results

### Behavioral results

#### Stress scores

Mental arithmetic successfully induced subjective stress. Stress ratings exhibited significant main effects of task and time point, as well as their interaction (F(1, 29) = 52.88, 18.47, and 20.72, all p’s < 0.001, respectively). Stress ratings were elevated at baseline before arithmetic blocks relative to passive viewing blocks (Fig. 2A) (t(29) = 4.13, p < 0.001), consistent with anticipatory stress. This elevation persisted post-task (t(29) = 7.49, p < 0.001) and post-rest (t(29) = 5.87, p < 0.001).

**Figure 1.**
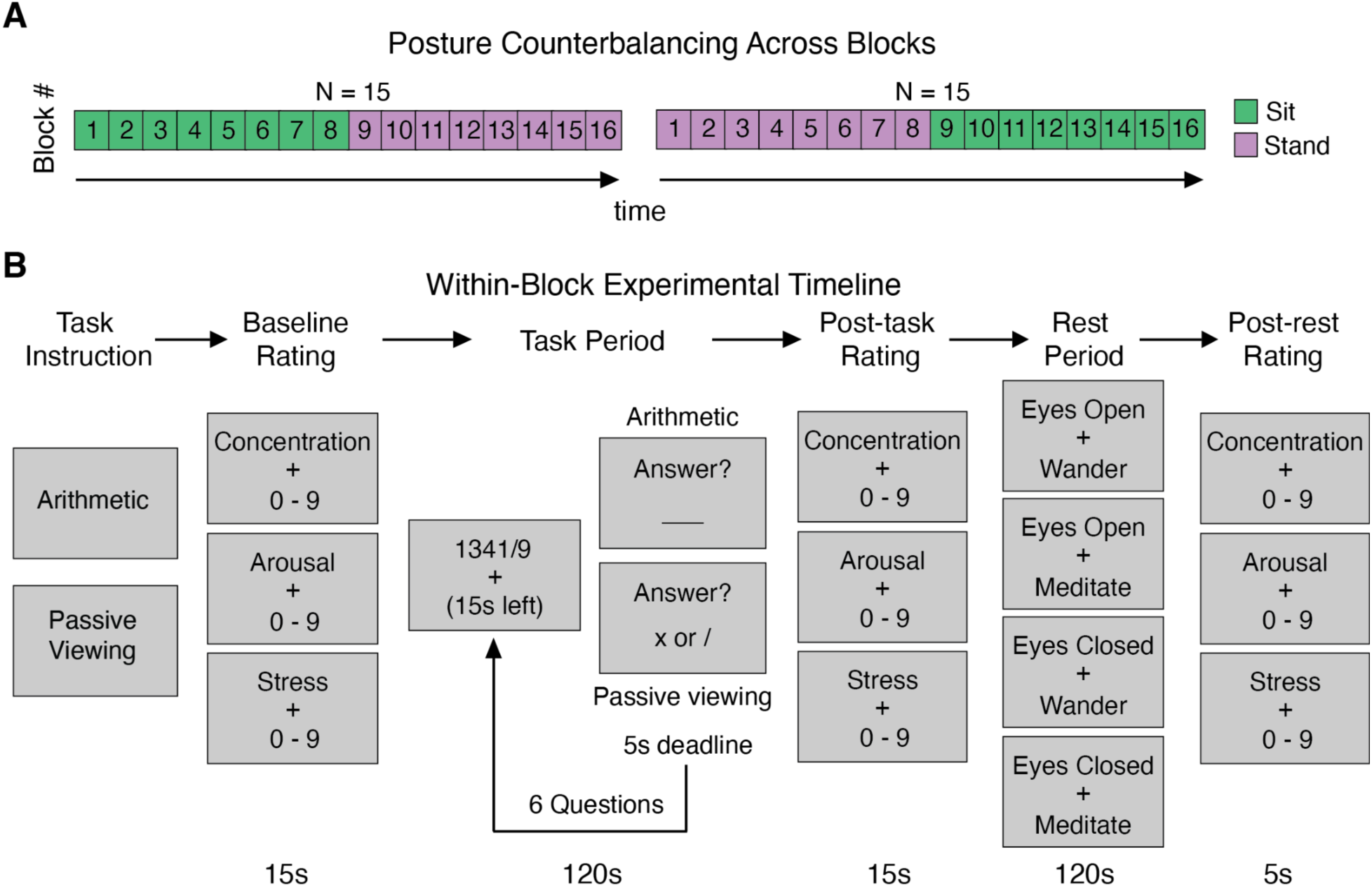
Experimental design for arithmetic stress induction and meditation recovery tasks. **(A)** Half of the participants (N = 15) completed eight blocks seated (green) followed by eight blocks standing (purple); the other half completed blocks in reverse posture order. **(B)** Regardless of posture, each experimental block began with an instruction screen indicating whether participants would perform mental arithmetic (math calculation) or passive viewing. Participants provided baseline subjective ratings of stress, arousal, and concentration on 0-9 Likert scales (5 seconds per rating). During the stress-induction phase, participants in the arithmetic condition solved six difficult math problems (multiplication or division). Participants had a 15-second time limit per problem and a 5-second response window for entering answers. In the passive viewing condition, participants viewed identical problems but only indicated whether each involved multiplication or division (x vs /), without performing calculations. Following the task phase, participants again rated their subjective states. This was followed by a 120-second rest period during which participants engaged in either mindful breathing meditation or mind-wandering, with eyes open or closed depending on condition assignment. A bell tone signaled the end of the rest period, after which participants completed final subjective ratings. All participants participated in block types crossing task conditions (arithmetic vs. passive viewing) and rest conditions (mind-wandering vs. meditation, eyes open vs. closed), and block order was counterbalanced across participants.

**Figure 2.**
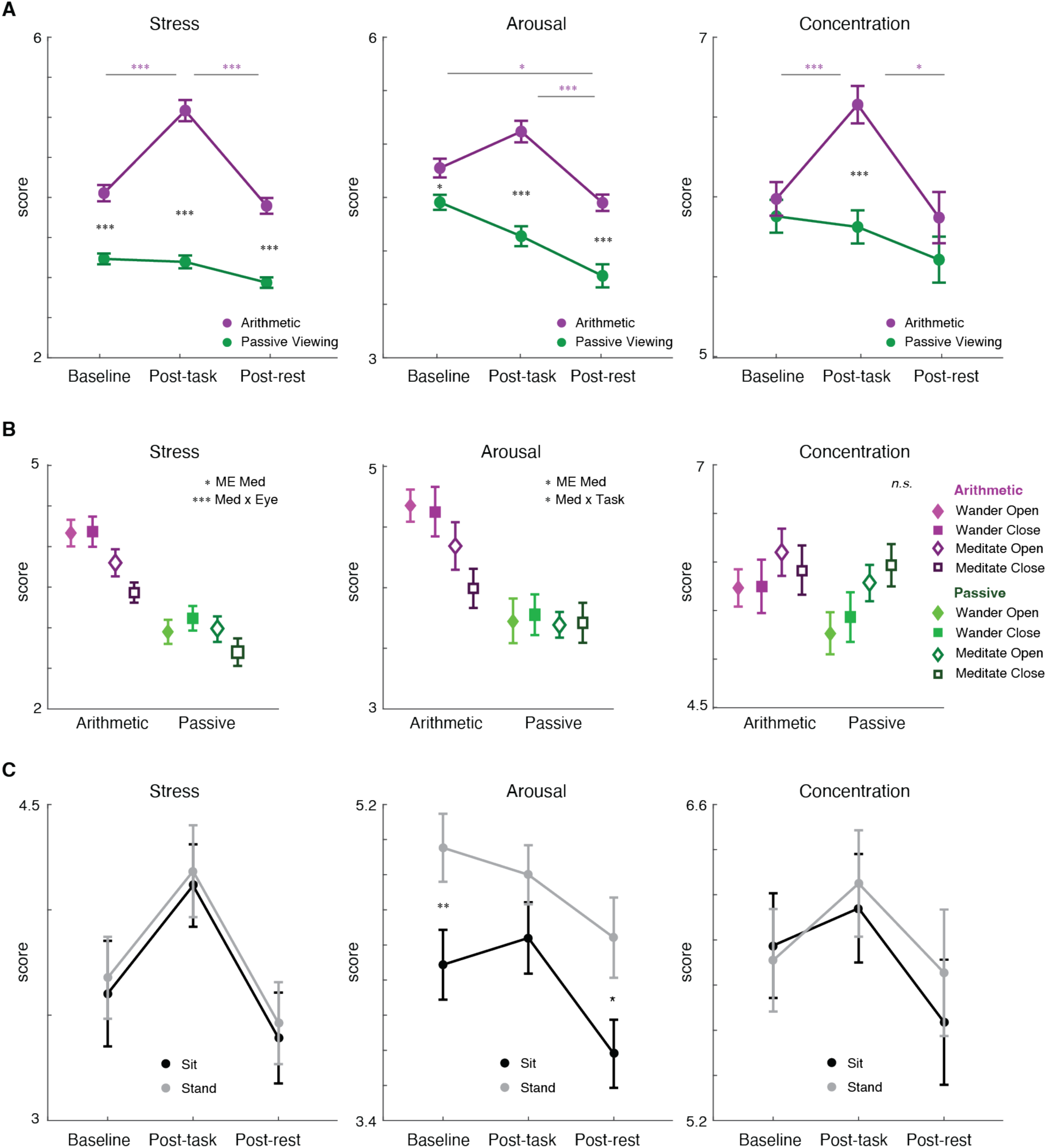
Mental arithmetic induces robust subjective stress that is reduced by mindful breathing, particularly with eyes closed. **(A)** Time course of subjective stress, arousal, and concentration ratings across baseline, post-task, and post-rest periods. Mental arithmetic (purple) significantly elevated all three measures relative to passive viewing (green), confirming successful stress induction. All measures decreased during the subsequent rest period, with stress and arousal falling below baseline levels. **(B)** Post-rest subjective ratings as a function of task context (arithmetic vs. passive viewing) and recovery condition. Eyes-closed mindful breathing produced greater stress reduction than eyes-open mindful breathing, particularly following arithmetic stress. Arousal showed a similar pattern, while concentration was not significantly modulated by meditation style or eye state. Purple symbols indicate arithmetic blocks, green symbols indicate passive viewing blocks; open symbols denote mindful breathing meditation, closed symbols represent mind wandering; squares denote closed eye conditions, diamonds denote open eye conditions. **(C)** Posture effects on subjective ratings. Standing (gray) increased subjective arousal relative to sitting (black) across all time points, but did not significantly affect stress or concentration ratings. Error bars represent within-subject SEM. * p < 0.05; ** p < 0.01; *** p < 0.001. ME = main effect; × = interaction.

Arithmetic increased stress from baseline (t(29) = 4.76, p < 0.001), whereas passive viewing produced no significant change (t(29) = –0.28, p = 0.7779) (Fig. 2A). The rest period reduced stress induced by arithmetic (t(29) = –5.65, p < 0.001) to baseline levels (t(29) = –1.06, p = 0.2982). Stress scores dropped below baseline after rest with passive viewing (t(29) = –2.77, p < 0.001) (Fig. 2A). Post-rest stress exhibited a main effect of meditation style (mindful breathing vs. mind-wandering) (F(1, 29) = 6.88, p = 0.0137) and an interaction between meditation and eye state (closed vs. open) (F(1, 29) = 8.61, p < 0.001) (Fig. 2A). This interaction reflected greater stress reduction for eyes closed versus eyes open meditation (t(29) = –2.93, p < 0.001), an effect was driven by the arithmetic condition (t(29) = –2.21, p = 0.0354) but absent following passive viewing (t(29) = –1.42, p = 0.1677) (Fig. 2A).

#### Arousal scores

Arousal ratings mirrored stress patterns, with significant main effects of task and time point, as well as their interaction (F(1, 29) = 28.66, 9.23, and 8.16, all p’s < 0.001, respectively). Elevated arousal ratings prior to arithmetic stress induction relative to passive viewing (t(29) = –2.06, p = 0.0488) (Fig. 2A) indicate anticipatory arousal consistent with the pattern observed for stress. Arithmetic marginally increased arousal (t(29) = 1.96, p = 0.0592), and rest reduced arousal ratings compared to the post-task (t(29) = – 4.25, p < 0.001) to a level below baseline (t(29) = –2.48, p = 0.0191) (Fig. 2A). Arousal during passive viewing continually decreased from baseline to post-task (t(29) = –2.54, p = 0.0168) (Fig. 2A) and subsequently post-rest (t(29) = –1.97, p = 0.0583) falling below baseline with rest (t(29) = –4.36, p < 0.001). Arithmetic effectively elevated arousal relative to passive viewing at post-task (t(29) = 5.93, p < 0.001) and post-rest time points (t(29) = 4.93, p < 0.001)(Fig. 2A).

Arousal showed a significant main effect of meditation (F(1, 29) = 6.25, p = 0.0183) and a significant interaction between meditation and task (F(1, 29) = 4.45, p = 0.0437), reflecting enhanced arousal reduction when mindfulness follows arithmetic compared to passive viewing (Fig. 2A)

Standing produced significantly higher arousal scores than sitting independently of mindfulness (F(1, 29) = 7.36, p = 0.0111) (Fig. 2B). This posture effect was evident at baseline (t(29) = 3.25, p = 0.0029) and post-rest (t(29) = 2.68, p = 0.0120), but not post-task time points (t(29) = 1.57, p = 0.1270). In contrast, posture had no significant effect on subjective stress (t(29) = 0.12, p = 0.7353) or concentration scores (t(29) = 0.12, p = 0.7265) (Fig. 2B), indicating posture specifically influences subjective arousal.

#### Concentration scores

Concentration showed significant main effects of task (arithmetic vs. passive viewing) and interaction between task and time (baseline, post-task vs. post-rest) (F(1, 29)’s = 8.67 and 7.04, p’s = 0.0063 and 0.0011, respectively). Unlike stress and arousal, baseline concentration levels between arithmetic and passive viewing did not differ (t(29)= 0.76, p = 0.4534). Concentration scores increased following arithmetic compared to baseline (t(29)= 3.81, p < 0.001) and decreased after the rest period (t(29) = −2.72, p = 0.0110), returning to baseline (t(29) = −0.48, p = 0.6362). Concentration under passive viewing remained at baseline throughout the entire block (t(29)’s=-0.89 to –1.19, p = 0.2437-0.3802) (Fig. 2A).

Unlike stress and arousal, post-rest concentration scores were similar after arithmetic and passive viewing but not at the post-rest time point (t(29) = 1.70, p = 0.1005), suggesting the increase in concentration (post-task time point (t(29) = 3.90, p < 0.001) was transient. Concentration scores after rest were not impacted by meditation (F(1, 29) = 2.58, p = 0.1188), eye state (F(1, 29) = 0.10, p = 0.7490), or their interaction (F(1, 29) = 0.33, p = 0.5693) (Fig. 2A).

### Neural Oscillatory Signatures of Stress

#### Scalp EEG

Mental arithmetic elicited four canonical oscillatory responses relative to passive viewing (Fig. 3). First, frontal-midline theta power (3-7 Hz) increased significantly (F(1, 29) = 15.93, p < 0.001), consistent with engagement of executive control, working memory maintenance, and error monitoring processes (Cavanagh & Frank, 2014; Gärtner et al., 2015; Hsieh and Ranganath, 2014; Itthipuripat et al., 2013; Jensen and Tesche, 2002; Sauseng et al., 2010). Second, posterior alpha power (10-12Hz) was suppressed during arithmetic (F(1, 29)’s = 14.31-22.57, p’s < 0.001 across left, mid, right posterior regions), reflecting attentional engagement and increased task demands (Banerjee et al., 2011; Bonnefond & Jensen, 2025; Foxe & Snyder, 2011; Itthipuripat et al., 2019, 2023; Jensen & Mazaheri, 2010; Klimesch et al., 1998, 2007, 2012; Nelli et al., 2017; Sookprao, 2024; Strauß et al., 2014; Woodman et al., 2022). Third, arithmetic suppressed low-beta power (15-20Hz) in central channels compared to passive viewing (F(1, 29)’s = 15.33-21.84, p’s < 0.001, p’s < 0.001 across left and right central electrodes), consistent with enhanced sensorimotor engagement (Kilavik et al., 2013; Meirovitch et al., 2015; Pfurtscheller & Lopes da Silva, 1999). Finally, posterior high-beta (25-32 Hz) and low gamma (35-48 Hz) power increased substantially (F(1, 29)’s = 7.50-18.19 with p’s = 0.0002-0.0105) with no significant modulation at other scalp locations (F(1, 29)’s = 0.00–3.96, p’s = 0.0562–0.9846). This focal posterior enhancement likely reflects increased cortical excitatory drive during sustained visual and attentional processing (Ahmad et al., 2022; Buzsáki & Wang, 2012; Whittington et al., 2000; Yokoyama et al., 2025). Taken together, these four spectral signatures confirm that mental arithmetic successfully induced cognitive stress and engaged attentional control mechanisms.

**Figure 3.**
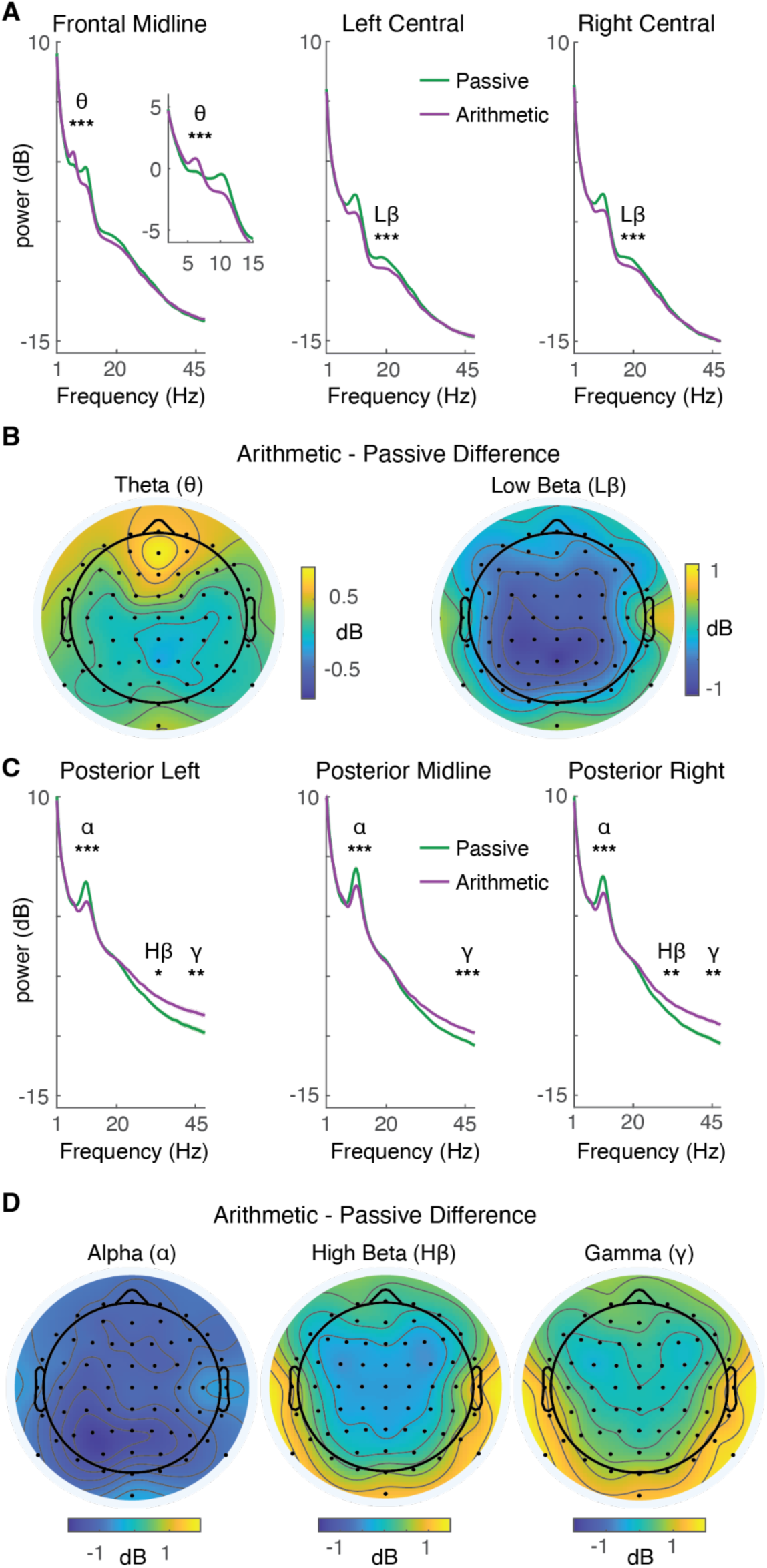
Mental arithmetic elicits canonical stress-related oscillatory signatures in scalp EEG. **(A)** Mental arithmetic (purple) increased midline frontal theta power (3-7 Hz) relative to passive viewing (green), consistent with enhanced cognitive control and working memory demands. Inset shows theta band average. Mental arithmetic also suppressed low-beta power (15-20 Hz) over bilateral central electrodes, reflecting enhanced motor preparation and response processes. **(B)** Topographic distributions of arithmetic minus passive viewing theta and low-beta effects. Theta enhancement was maximal over midline frontal electrodes. Low-beta suppression was broadly distributed across central regions. **(C)** Mental arithmetic (purple) suppressed posterior occipital alpha power (10-12 Hz) bilaterally, indicating increased attentional engagement. Mental arithmetic also elevated high-beta (25-32 Hz) and gamma (35-48 Hz) power over posterior occipital regions, reflecting increased excitatory drive and heightened visual processing demands. **(D)** Topographic distributions of arithmetic minus passive viewing alpha, high-beta, and gamma effects. Alpha suppression and high-beta/gamma enhancement were maximal over posterior occipital scalp. Black dots indicate electrode locations. Shaded regions represent within-subject SEM. *, p < 0.05; **, p < 0.01; ***, p < 0.001.

Posture modulated arousal-related activity independent of cognitive stress. Standing significantly elevated high-beta and gamma power across posterior regions compared to sitting (F(1, 29)’s = 7.77-24.07, 19.46 with p’s <0.001 to 0.0093) (Fig. 4). This paralleled increased subjective arousal ratings during standing (Fig. 2) and likely reflects heightened cortical arousal or visual vigilance associated with upright posture rather than stress per se. Critically, stress-induced oscillatory effects reported above were consistent across sitting and standing (significant main effects of task as reported above without task-by-posture interactions; F(1, 29)’s = 0.04-3.53, p’s = 0.0702-0.8456), demonstrating the posture-invariance of EEG stress signatures. Importantly, increased high-frequency activity during standing could potentially be confounded by muscular activity arising from postural muscle engagement, particularly in the neck and shoulder regions. To mitigate this possibility, ICA was applied to remove components reflecting muscle and movement artifacts prior to spectral analyses. Moreover, the observed effects were spatially concentrated over posterior occipital scalp regions rather than lateral electrode sites typically associated with neck muscle activity, arguing against a purely muscle-related origin.

**Figure 4.**
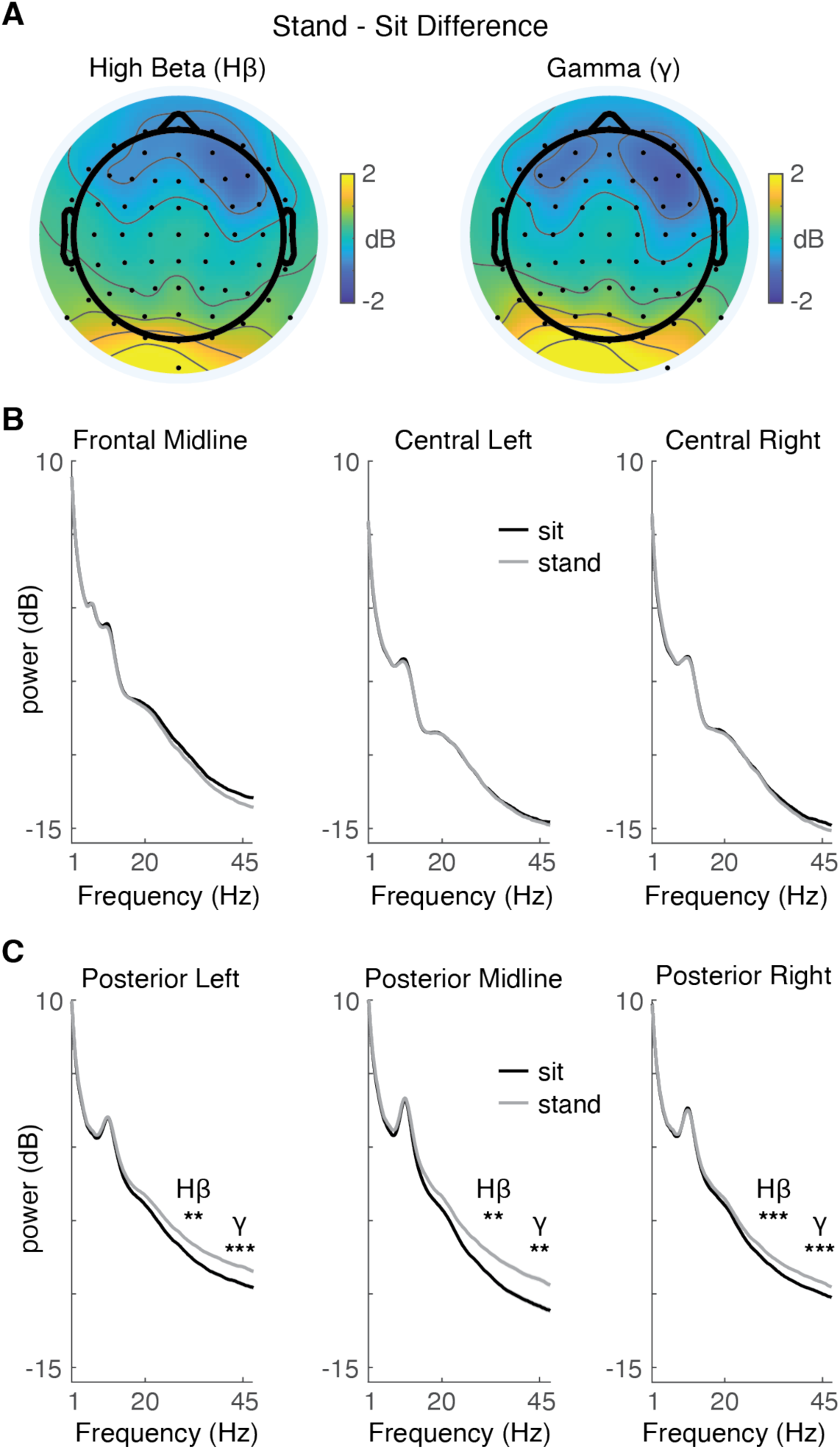
Standing posture increases posterior high-frequency activity. **(A)** Topographic distributions showing enhanced high-beta (left) and gamma (right) power during standing relative to sitting, concentrated over posterior occipital regions. This pattern paralleled increased subjective arousal during standing (Figure 2C) and likely reflects heightened cortical arousal or visual vigilance associated with upright posture. **(B)** Power spectra from midline frontal and bilateral central electrode groups showing no significant posture effects at these locations. **(C)** Power spectra from bilateral posterior occipital electrode groups showing robust high-beta and gamma enhancement during standing (light gray) relative to sitting (black). Critically, stress-induced oscillatory effects remained consistent across postures, demonstrating posture-invariance of EEG stress signatures. Shaded regions represent within-subject SEM. ** p < 0.01; *** p < 0.001.

#### Around-ear EEG

Around-ear electrodes captured two of the four canonical stress-related oscillatory responses observed in full-scalp recordings (Fig. 5A). Alpha suppression during mental arithmetic was reliably detected across all around-ear contacts (F(1, 29)’s = 7.38-13.54, p’s < 0.001, surviving Holm-Bonferroni correction), demonstrating robust propagation of this signature to peripheral sites. Gamma power increased significantly during arithmetic, most pronounced at right frontal and temple contacts (F(1, 29)’s =7.69-8.09, p’s = 0.0081-0.0096), demonstrating sensitivity to stress-related high-frequency activity. These stress-related neural dynamics were consistent across sitting and standing postures (significant main effects of task without task-by-posture interactions: F(1, 29)’s 0.0119-2.4996, p’s = 0.1247 – 0.9140), confirming that around-ear EEG can reliably capture cognitive stress signatures independent of body position.

**Figure 5.**
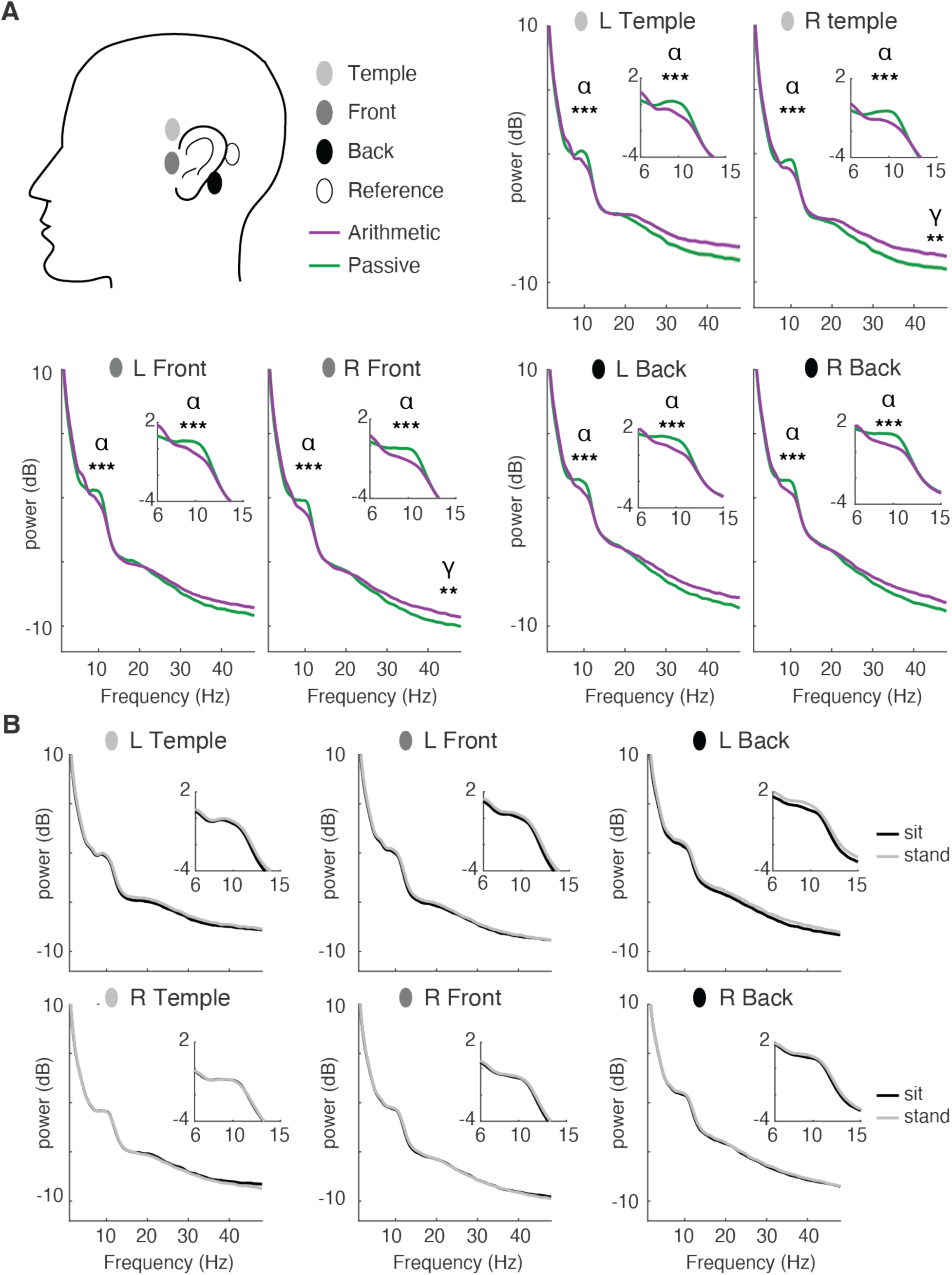
Around-ear EEG captures stress-related alpha suppression and gamma enhancement. **(A)** Schematic showing around-ear electrode locations (left). Power spectra from six around-ear electrode contacts during mental arithmetic (purple) versus passive viewing (green). Mental arithmetic suppressed alpha power (10-12 Hz) across all electrode locations, demonstrating robust propagation of this signature to peripheral sites. Mental arithmetic also elevated gamma power (35-48 Hz), with effects most pronounced at right frontal and bilateral temple contacts. Insets show frequency bands of interest. **(B)** Posture effects on around-ear EEG. Unlike scalp recordings, around-ear electrodes showed no significant high-beta or gamma modulation by posture (sitting, black; standing, gray). This spatial selectivity indicates that around-ear contacts preferentially capture stress-related temporal lobe activity while showing reduced sensitivity to posterior arousal-related signals. Shaded regions represent within-subject SEM. ** p < 0.01; *** p < 0.001.

Critically, around-ear recordings showed no significant posture-related modulation of high-beta or gamma power (F(1, 29)’s = 0.0274-2.0962, p’s = 0.1584-0.8697) (Fig. 5B), contrasting with posterior arousal-related increases. This spatial specificity indicates that around-ear electrodes selectively capture stress-related neural activity while showing limited sensitivity to arousal-related signatures. Together, these findings establish the feasibility of around-ear EEG for naturalistic assessments of cognitive states such as stress in seated and upright postures.

### Neural Signatures During Stress Recovery

#### Sustained stress-related responses during rest in Scalp EEG

We assessed neural recovery by comparing rest-period oscillatory activity following arithmetic versus passive viewing across meditation and eye-state conditions. Unlike stress induction, which modulated multiple frequency bands across widespread scalp regions, the rest period exhibited selective high-frequency modulation, with attenuated or reversed lower-frequency effects.

Bilateral posterior and temporal regions exhibited sustained high-beta and gamma power elevations following mental arithmetic compared with passive viewing (F(1, 29)’s = 5.04–9.03, p’s = 0.0326–0.0054; Fig. 6). In contrast, midline theta and posterior alpha power showed attenuated, reversed modulation patterns relative to those observed during stress-induction. Arithmetic decreased midline theta power (F(1, 29) = 4.80, p = 0.0367) and increased bilateral posterior alpha power (F(1, 29)’s = 4.43–4.51, p’s = 0.0423–0.0440) relative to passive viewing. Central low-beta activity showed no significant differences between task contexts (F(1, 29)’s = 1.40–3.55, p’s = 0.0694–0.2459).

**Figure 6.**
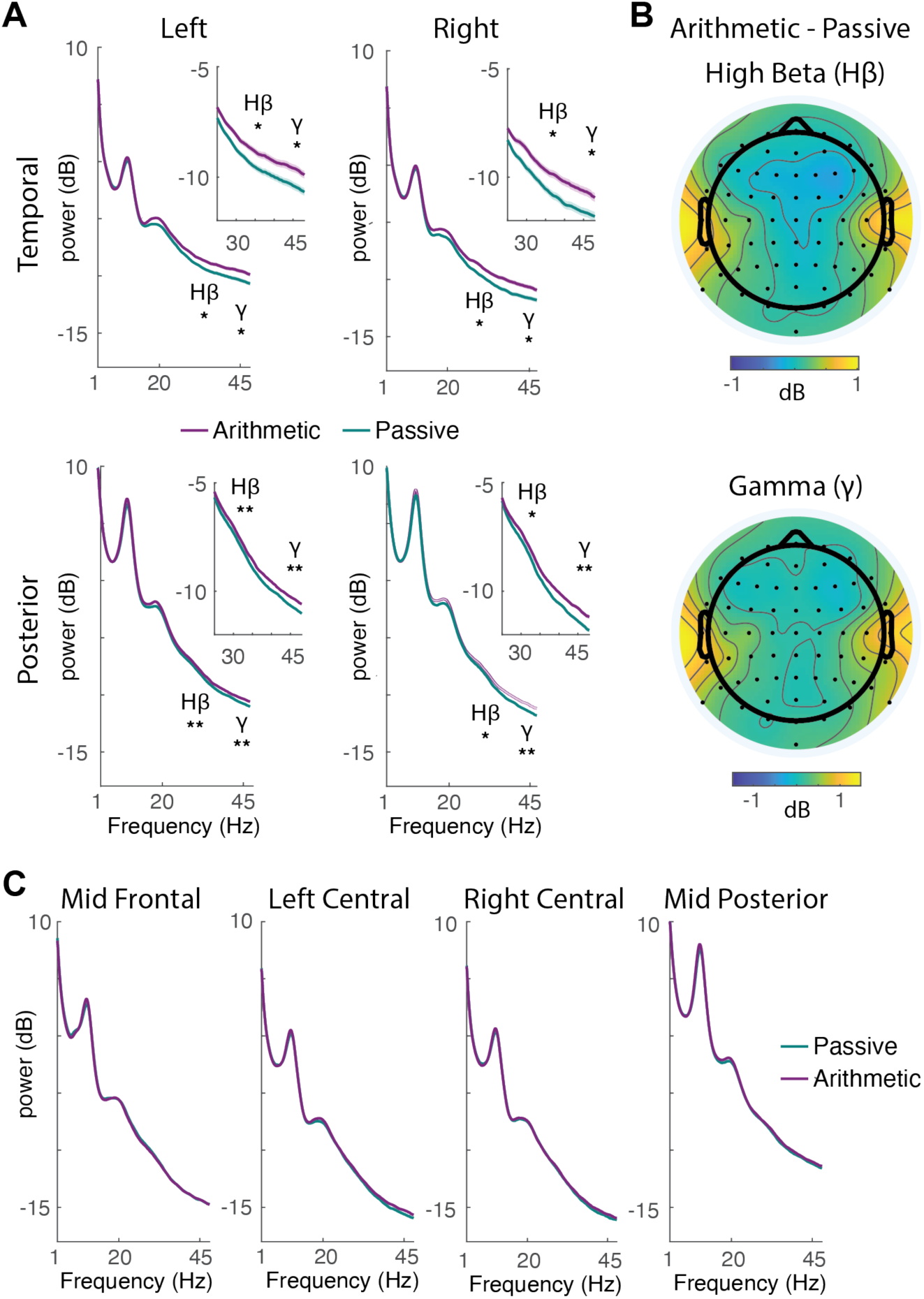
Stress-related high-frequency activity persists during the post-task rest period. **(A)** Power spectra from bilateral temporal (top) and posterior occipital (bottom) scalp electrodes during rest periods following mental arithmetic (purple) versus passive viewing (green/cyan). High-beta (25-32 Hz) and gamma (35-48 Hz) power remained elevated following arithmetic stress, indicating sustained neural excitation outlasting the acute stressor. **(B)** Topographic distributions of high-beta and gamma power differences (arithmetic minus passive viewing) during rest. Elevated high-frequency activity was concentrated over bilateral temporal and posterior occipital regions. **(C)** Power spectra from midline frontal, bilateral central, and mid-posterior occipital electrode groups. Unlike the stress-induction period, midline frontal theta decreased following arithmetic relative to passive viewing, and posterior alpha showed reversed modulation (increased rather than suppressed). Central low-beta showed no significant task-related differences during rest. Black dots on topographic maps indicate electrode locations. Shaded regions represent within-subject SEM. *, p < 0.05; **, p < 0.01.

Sustained high-frequency elevation following stress induction suggests persistent neural excitation outlasting the acute stressor, consistent with incomplete neural recovery. Notably, high beta and gamma band responses remained consistent across sitting and standing, with no significant interaction between task context and posture (F(1,29)’s = 0.00–0.86, p = 0.3610–0.9576). However, a localized increase in high-beta and gamma power was observed in posterior occipital regions during standing compared with sitting (F(1,29)’s =9.78–18.37, p’s = 0.0002–0.0040). This pattern mirrors the stress-induction phase results and likely reflects visuospatial processing and postural control demands during upright stance.

#### Lingering stress-related responses in around-ear EEG

Around-ear recordings corroborated scalp EEG findings. Bilateral electrodes positioned anterior to the temples showed sustained elevations in high-beta and gamma power following mental arithmetic (F(1, 29)’s = 9.19–14.11, p’s = 0.0008–0.0051), all surviving Holm–Bonferroni correction for comparisons across the six around-ear contacts. In contrast, low-frequency EEG activity did not show significant modulation after correction (Theta and Alpha: F(1, 29)’s = 0.64–6.38, p’s = 0.0173–0.4318, corrected threshold = 0.0083). This frequency-specific pattern supports high-beta and gamma activity as markers of prolonged stress-related neural excitation, whereas theta and alpha modulations observed during task performance likely reflect active attentional and cognitive operations rather than enduring stress responses.

**Figure 7.**
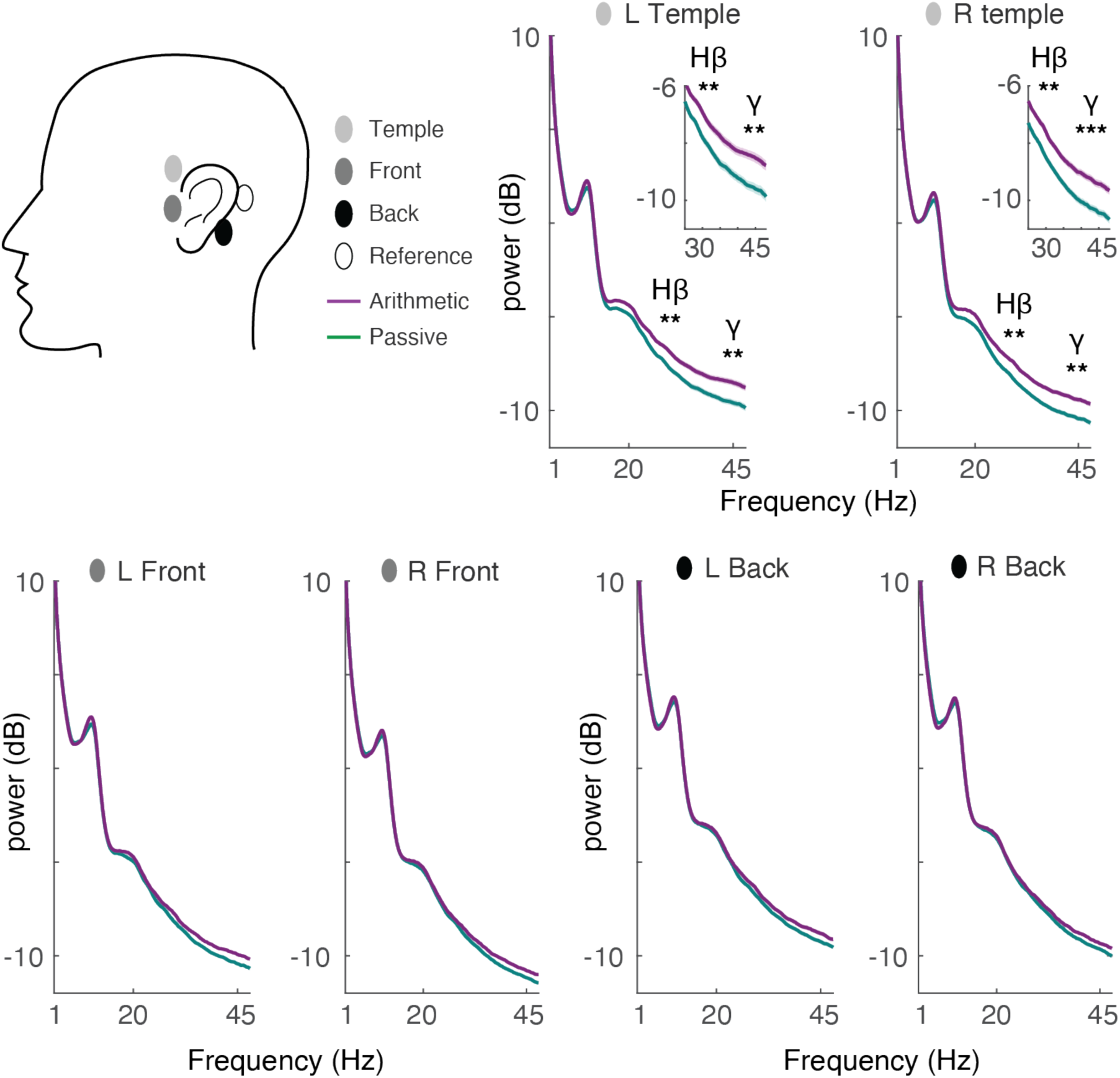
Around-ear EEG detects sustained stress-related high-frequency activity during rest. Schematic showing around-ear electrode locations (left). Power spectra from six around-ear electrode contacts during rest periods following mental arithmetic (purple) versus passive viewing (green). Consistent with scalp recordings, around-ear electrodes positioned at bilateral temple locations showed sustained elevation of high-beta (25-32 Hz) and gamma (35-48 Hz) power following arithmetic stress. These effects survived Holm-Bonferroni correction for multiple comparisons across the six around-ear contacts. In contrast to high-frequency activity, theta and alpha power showed no significant task-related modulation during rest after correction, indicating these frequency bands reflect active cognitive operations during task performance rather than enduring stress responses. Shaded regions represent within-subject SEM. ** p < 0.01; *** p < 0.001.

#### Context-Dependent Modulation by Meditation and Eye State

We next examined how meditation style (mindful breathing vs mind wandering) and eye state (open vs closed) shaped neural activity during stress recovery. Participants completed four resting conditions following mental arithmetic or passive-viewing: eyes open mindful breathing, eyes closed mindful breathing, eyes open mind-wandering, and eyes closed mind-wandering (Fig. 1; see Materials & Methods). We analyzed rest-period EEG in two temporal windows to capture dynamic effects. The early window (0–20 s) captured rapid neural transitions following rest onset, while the late window (20–120 s) reflected sustained recovery dynamics (data collapsed across 20 s bins given stable spectral patterns). This time-resolved approach distinguished transient effects emerging at rest onset from sustained influences of meditation and sensory state on stress-related neural signatures.

Eye closure robustly increased posterior alpha power independent of meditation style (F(1, 29) = 12.11, p = 0.0016), and time window (eye state × time: F(1, 29) = 0.03, p = 0.8536). This alpha enhancement extended across all scalp regions (F(1, 29)’s = 54.85-69.62, p’s < 0.001, all surviving Holm-Bonferroni correction across eight scalp electrode groups) and all around-ear electrodes (F(1, 29)’s = 31.48-43.46, p’s < 0.001, all surviving Holm-Bonferroni correction across six around-ear EEG contacts). Meditation condition (mindful breathing versus mind wandering) did not significantly modulate alpha band activity (F(1, 29) = 3.36, p = 0.0771) and did not interact with eye state (F(1, 29) = 0.56, p = 0.4590), task context (F(1, 29) = 0.16, p = 0.6938), or time window (F(1, 29) = 0.03, p = 0.8636).

In contrast to alpha activity, we observed significant four-way interactions among meditation style, eye state, task context (post-arithmetic vs. post-passive viewing), and time window (0–20s vs. 20–120s) on gamma activity recorded over temporal scalp regions and temple contacts of around-ear EEG (F(1, 29)’s = 10.79 and 5.91, p’s = 0.0207 and 0.0215, respectively). These effects reflected distinct modulatory patterns of gamma activity across the early and late time windows.

During the early window (0–20s), we observed significant three-way interactions between meditation condition, eye state, and task on gamma band activity (F(1, 29)’s = 15.36 and 9.77, p’s = 0.0005 and 0.004 for temporal scalp and temple around-ear EEG contacts, respectively). These interactions indicated that eye closure differentially modulated meditation-related gamma reductions depending on prior task. Following arithmetic, eyes-closed mindful breathing suppressed gamma power more strongly than eyes-open (t(29)’s = 2.06 and 1.84, p’s = 0.0240 and 0.0380 for temporal scalp and temple around-ear EEG contacts respectively; one-tailed, based on direction of interaction). Following passive viewing, eyes-open mindful breathing reduced gamma power (t(29)’s = –3.79 and –2.61, p’s = 0.0003 and 0.0071 for temporal scalp and temple around-ear EEG contacts respectively; one-tailed, based on the known direction of the interaction), whereas eyes-closed mindful breathing produced no additional reduction.

Late (20-120s) gamma power decreased relative to the early period (F(1,29)’s = 47.84 and 53.90, p’s < 0.001 for temporal scalp and temple around-ear EEG contacts respectively), indicating progressive reduction of stress-related neural activity. During the late window, only eye state (F(1, 29)’s = 21.52 and 10.99, p’s =0.0001 and 0.0025) and task (F(1, 29)’s = 5.36 and 12.40, p’s = 0.0279 and 0.0014 for scalp and around-ear EEG, respectively) significantly modulated gamma power, with no impact of meditation (Main Effect: F(1, 29)’s = 0.00 and 0.07, p’s = 0.9490 and 0.7998; Meditation x eye state x task interaction: F(1, 29)’s =2.73 and 2.15, p’s = 0.1093 and 0.1533 for scalp and around-ear EEG, respectively). This effect was primarily driven by reduced gamma power with eyes-closed compared with eyes-open. Together, these findings indicate that meditation-related differences are most pronounced immediately following stress induction, whereas later in recovery, gamma activity becomes less sensitive to meditation practice and primarily reflects eye state. Importantly, convergent patterns across temporal scalp electrodes and corresponding around-ear temple contacts demonstrate that around-ear EEG reliably tracks stress– and meditation-related high-frequency dynamics in temporal brain regions.

**Figure 8.**
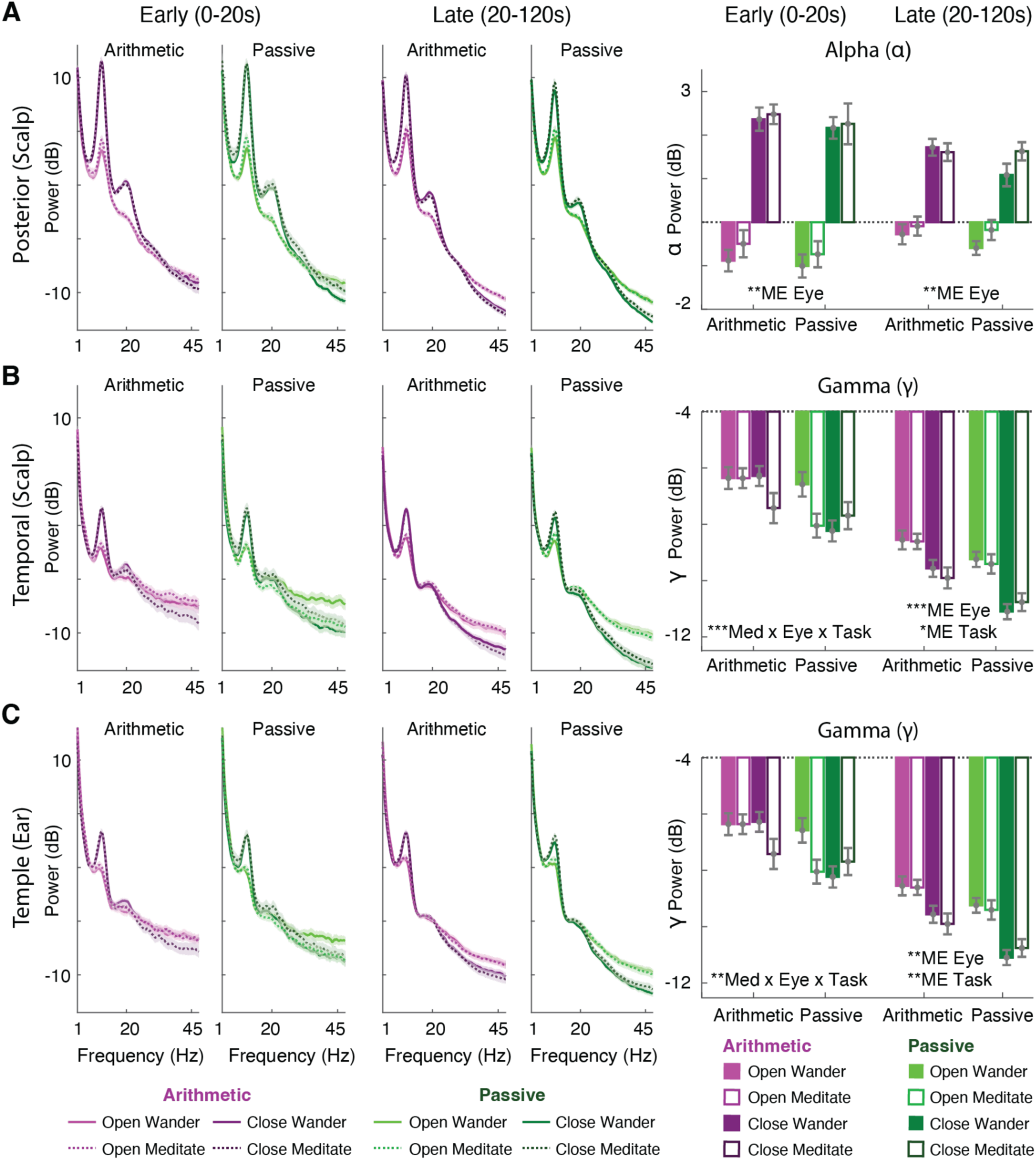
Mindful breathing dynamically modulates activity in a context-dependent manner. **(A)** Alpha power from posterior occipital scalp electrodes (left panels) and summary data (right panels) showing robust enhancement during eyes-closed conditions (filled bars) relative to eyes-open conditions (open bars), independent of meditation style. Purple bars indicate arithmetic blocks; green bars indicate passive viewing blocks. This effect was evident during both early (0-20 s) and late (20-120 s) rest windows, though it diminished over time. Topographic map shows maximal alpha enhancement over posterior regions. **(B)** Gamma power from bilateral temporal scalp electrodes (left panels) and summary data (right panels) showing a significant three-way interaction between meditation style, eye state, and prior task context during the early (0-20s) rest window. Following arithmetic stress, eyes-closed mindful breathing (dark filled bars) produced maximal gamma suppression. Following passive viewing, eyes-open mindful breathing (light open bars) reduced gamma relative to eyes-open mind-wandering, while eyes-closed meditation showed no additional benefit. During the late window (20-120s), meditation-specific effects diminished, with gamma modulation primarily reflecting main effects of eye state and prior task. **(C)** Gamma power from bilateral temple contacts of around-ear EEG showing convergent patterns with temporal scalp recordings, validating around-ear detection of meditation-related high-frequency dynamics. Color coding: purple/magenta indicate arithmetic blocks; dark green/light green indicate passive viewing blocks. Meditation conditions shown by darker shades; mind-wandering shown by lighter shades. Solid bars indicate eyes-closed; open bars indicate eyes-open. Shaded regions and error bars represent within-subject SEM. * p < 0.05; ** p < 0.01; *** p < 0.001. ME = main effect; × = interaction.

## Discussion

We demonstrate that periauricular EEG reliably captures neural signatures of stress and mindfulness-based recovery, establishing the viability of minimally invasive wearable systems for real-world mental health monitoring. Direct comparison of full-scalp and periauricular recordings revealed convergent evidence that mindful breathing rapidly modulates stress-related neural dynamics in a context-dependent manner, with efficacy determined by prior cognitive state and eye position. These findings bridge fundamental questions about meditation mechanisms with practical advances in accessible neurofeedback technology

### Context-Dependent Neural Regulation During Mindful Breathing

Mindful breathing reduced subjective stress (Kabat-Zinn, 1985; Segal et al., 2002), with maximal efficacy during eyes-closed practice. This finding suggests that optimal recovery requires both directed interoceptive attention and minimal exteroceptive interference. This pattern aligns with frameworks proposing that meditation enhances interoceptive awareness by attenuating exteroceptive processing (Farb et al., 2013; Fox et al., 2012; Gibson, et al., 2019; Josipovic et. al., 2013; Khalsa, et al., 2008; Tang et al., 2015). The selective enhancement of meditation efficacy by eye closure following stress, but not passive viewing, suggests adaptive recruitment of stronger regulatory interventions in response to internal dysregulation (Arch & Craske, 2006; Tang et al., 2015; Vago & Silbersweig, 2012).

Selective arousal elevation during standing, independent of stress or concentration, provides evidence for dissociable physiological and cognitive contributions to subjective state. This suggests posture may primarily affect autonomic tone and vigilance systems rather than cognitive load per se, consistent with embodied cognition frameworks linking body position to neural state (Cacciatore et al., 2024; Muehlhan et al., 2014; Nair et al., 2015; Takayama & Sekiya, 2023; see review Barsalou, 2008). Future work examining how physical and cognitive stressors interact during recovery could inform more nuanced intervention design.

### Oscillatory Mechanisms of Stress Induction and Recovery

Elevated temporal gamma activity persisted into the rest period, suggesting incomplete neural recovery despite subjective relief. This dissociation between subjective and neural stress markers raises critical questions about differential time courses of experiential versus neurophysiological recovery. While theta, alpha, and beta rapidly normalized during the recovery period, the selective persistence of gamma power implicates high-frequency oscillations as sensitive biomarkers of residual cortical excitation that may index individual differences in stress vulnerability and recovery trajectories.

Gamma suppression during eyes-closed meditation likely reflects convergent mechanisms. Eye closure reduces bottom-up sensory drive to visual cortex, decreasing baseline excitability (Barry et al., 2007; Berger, 1929; Cahn & Polich, 2006; Jensen & Mazaheri, 2010; Toscani et al., 2010). Simultaneously, directed attention to breathing engages top-down regulatory circuits suppressing task-irrelevant cortical activity (Jensen & Mazaheri 2010; Tang et al., 2015). The requirement for both factors suggests stress-induced cortical excitation necessitates strong regulatory intervention. This interpretation aligns with models whereby meditation training enhances cognitive control over distributed cortical networks (Cahn & Polich, 2006; Tang et al., 2015).

Meditation did not impact eye-closure-induced alpha enhancements reflecting established thalamocortical dynamics during reduced visual input (Barry et al., 2007; Klimesch et al., 2007). The gradual convergence between eyes-open and eyes-closed alpha may reflect natural relaxation processes over time or, alternatively, convergence of mind-wandering and meditative states during unstructured rest.

### Rapid Neural Modulation Within Seconds of Practice Onset

Meditation effects emerged within 20 seconds of practice onset, demonstrating rapid neural state modulation. This timescale contrasts longitudinal studies documenting gradual neural plasticity over weeks to months (Cahn et al., 2006; Hölzel et al., 2011; Itthipuripat et al., 2017; Saggar et al., 2012; Skwara et al., 2012; Tang et al., 2015; Yoshida et al., 2020). Acute practice may provide immediate relief through state regulation, while sustained training may drive structural or functional plasticity strengthening regulatory capacity.

The attenuation of meditation-specific effects during later rest periods could reflect non-mutually exclusive processes. First, all conditions may converge toward a common resting state as acute stress dissipates naturally over time. Second, brief meditation instructions may produce time-limited effects without sustained practice. Third, individual differences in practice adherence during the two-minute period may introduce variability that obscures condition differences. Future studies using continuous subjective reports or behavioral indices of meditation engagement could disambiguate these possibilities.

### Wearable Around-Ear EEG Validation for Stress Monitoring

Successful detection of stress and meditation signatures using around-ear electrodes represents a critical methodological advance. Previous studies validated around-ear EEG for evoked potentials and tonic oscillatory rhythms (Athavipach et al., 2019; Bleichner & Debener, 2017; Bleichner et al., 2015; Debener et al, 2015; Hemakom et al., 2024; Israsena & Pan-ngum, 2022; Kaongoen et al., 2021; Sanguantrakul et al., 2024;, see reviews Choi et al., 2023; Correia et al., 2024; da Silva Souto et al., 2022; Geirnaert et al., 2025; Hans et al., 2022; Kaongoen et al., 2023; Knierim et al., 2023; Mai et al., 2025; Petrossian et al., 2023), our findings extend this validation to dynamic state changes relevant for mental health applications. The observed spatial selectivity, robust temporal gamma detection but limited posterior sensitivity, demonstrates that around-ear electrode positioning can be strategically optimized for target applications.

Convergent temporal gamma patterns across around-ear and full-scalp recordings validate high-frequency temporal activity as a reliable stress biomarker. Posture-invariance across sitting and standing conditions further strengthens translational potential by demonstrating robustness across physical contexts. Limited around-ear sensitivity to posterior arousal-related gamma modulation may actually enhance stress detection specificity by excluding confounding arousal signals

### Limitations and Future Directions

Several limitations warrant consideration. Our sample included participants with prior meditation experience, potentially enhancing responsivity to brief instructions. Generalization to meditation-naive individuals requires systematic investigation. We examined only acute cognitive stress via mental arithmetic. Extension to social-evaluative stress, emotional challenges, and chronic stress would strengthen ecological validity across stressor types. Brief rest periods (2 minutes) may have precluded detection of sustained effects emerging during longer meditation sessions or accumulating across repeated practice bouts. Around-ear electrodes captured temporal activity but provided limited access to frontal and posterior sources. Hybrid systems integrating around-ear placement with strategic portable sensors could achieve broader coverage while maintaining practical wearability. Finally, we did not assess whether baseline neural states predict recovery trajectories or intervention outcomes. Longitudinal studies linking acute oscillatory biomarkers to long-term mental health outcomes could establish prognostic value.

These findings establish groundwork for adaptive neurofeedback systems tailoring interventions to individual neural states. Closed-loop systems monitoring temporal high-frequency activity could provide context-dependent guidance: prompting eyes-closed meditation when stress-related excitation is elevated, and eyes-open practice during lower-stress states. Rapid meditation effects within 20 seconds suggest that brief practice bouts integrated into daily routines could provide meaningful clinical benefit.

However, individual differences in meditation experience, baseline neural dynamics, and stress responsivity likely modulate efficacy, necessitating personalization algorithms that learn individual signatures and optimize interventions across diverse populations.

## Conclusions

Simultaneous full-scalp and periauricular EEG recordings during stress induction and mindfulness-based recovery demonstrate that wearable neurotechnology can reliably track stress-related neural dynamics. Mindful breathing modulates temporal high-frequency activity in a context-dependent manner, with maximal gamma suppression during eyes-closed practice following acute stress. Rapid neural modulation within 20 seconds of practice onset, combined with reliable periauricular detection during naturalistic movement, establishes feasibility for real-world neurofeedback applications. These findings bridge contemplative neuroscience with accessible technology, enabling personalized mental health interventions grounded in direct neural state monitoring.

## Data and code availability statement

The data and analysis code supporting the findings of this study are available from the corresponding author upon reasonable request, subject to institutional and ethical approval due to participant privacy considerations.

## Author Contributions and Acknowledgement

SN, CM, and Sirawaj Itthipuripat (SI1) conceived and designed the study. WW, TP, AS, and Suparach Intarasopa (SI2) collected the data. SN, PS, WW, RP, and SI1 analyzed the data. SN, PS, SK, MH, BG, and SI1 interpreted the results. SN, PS, and SI1 drafted the original manuscript. BG and SI1 supervised the project and critically revised the manuscript. CM, PS, and SI1 secured funding for the project. This project was supported by Brainspoke Lab and Alphie Holding Pte. Ltd., Singapore, provided to the Neuroscience Center for Research and Innovation, KMUTT; the Mid-Career Researcher Development Grant (2026–2028) from the National Research Council of Thailand (NRCT) awarded to SI1; the Frontier Research Fund awarded to the Neuroscience Center for Research and Innovation, KMUTT, by the Thailand Science Research and Innovation (TSRI) agency (to SI1); and the National Science Research and Innovation Fund (NSRI; 2026) through the Program Management Unit for Technology and Innovation for Future Industries (PMU-B), awarded to SI1 and PS.

